# Role of nitric oxide in improving seed germination and alleviation of copper-induced photosynthetic inhibition in Indian mustard

**DOI:** 10.1101/865162

**Authors:** Bilal A. Rather, Iqbal R. Mir, Asim Masood, Naser A. Anjum, Nafees A. Khan

## Abstract

Heavy metal stress limits crop production through its effects on seed germination and photosynthesis. Nitric oxide (NO), a versatile signaling molecule, plays a significant role in heavy metal stress tolerance. In the present investigation, the efficacy of NO application in the alleviation of copper (Cu) induced adverse impact on seed germination and photosynthesis of mustard plant (*Brassica juncea L*.) was evaluated. Pretreatment with NO donor sodium nitroprusside (SNP), significantly improved seed germination and alleviated Cu-accrued oxidative stress in *B. juncea* seeds. However, in the absence of NO, Cu showed a higher reduction in seed germination rate. Further, NO modulated the activities of antioxidant enzymes and sustained the lower level of lipid peroxidation by reducing H_2_O_2,_ and thiobarbituric acid reactive substances (TBARS), thereby elevated the antioxidative capacity in Cu-exposed seeds. Seeds pretreated with NO also retained higher amylase activities for the proper seed germination when compared with control. NO mitigated Cu toxicity through an improved antioxidant system, and reducing Cu-induced accumulation of reactive oxygen species (ROS), reduction in lipid peroxidation improving photosynthetic efficiency and growth of the mustard plant. It may concluded that NO improved amylase activity, modulated activity of antioxidant enzymes, and enhanced the germination rate seeds under Cu stress, thereby improved photosynthesis and growth.

## Introduction

Heavy metal pollution has been well-known since long time; however, its continuous use becomes hazardous in respect of agricultural and human health, mostly in lesser developed countries [1, 2]. In this perspective, Cu became apparent as a severe pollutant because of its extensive use in industries and as a pesticide in agriculture [3, 4]. The metal Cu performs a remarkable array of functions in living organisms that are essential for life. Proteins that contain Cu as cofactor perform various biochemical processes which are involved in plant growth and development, and protective mechanisms [5]. For plant life, Cu is widely recognized as an essential microelement [4]. Usually, the optimum levels of Cu in leaves are 10 µg g^-1^ dry mass [6] and the acute level of toxicity found for most of the crop plants are slightly higher than 20-30 µg g^-1^ dry mass [7]. Phytotoxicity of Cu in leaves affects photosynthesis adversely by targeting thylakoids; particularly PSII has been identified to be the main target [8]. Excessive accumulation of Cu dissipates the electron transport in PS I and PS II, inhibits photophosphorylation and alters membrane integrity [9, 10]. Also, high concentrations of Cu have been extensively reported to induce oxidative stress resulting in disruption of structure of biological molecules and membranes by increasing lipid peroxidation [11, 12]. Moreover, excess Cu alters the proper functioning of mineral uptake[13, 14]. Assay of seed germination is an essential procedure to investigate the Cu toxicity in various crop plants, while lower concentrations of Cu seemed to be significant for seed germination. Some of the known influence of Cu toxicity on bean seeds are (1) alteration of seed germination metabolism and (2) interference in the pathway of ubiquitin-proteasome mechanism to oxidatively-damaged proteins [15]. Cu at higher concentrations seized rate of radical elongation, especially in *Alyssum montanum* and *Thlaspio chroleucum* [16]. The special effects of Cu on seed germination are well documented [17–20].

To improve plants adaptability, phytohormones could serve as a potential tool under abiotic stress conditions. In particular, nitric oxide (NO) has been found to play a critical role in plant defense reactions and therefore, could serve as a promising plant hormone under various plant abiotic stresses [21, 22]. There is sufficient evidence of literature that substantiates wide range of roles of NO in plant physiological processes, such as germination, photosynthesis, root development, flowering, stomatal closure, pollen tube growth regulation, senescence, fruit ripening and defensive mechanism under various environmental stresses [23, 24]. In plants, the cell-protective role of NO has openly been tested with DNA, proteins, lipids and chlorophyll [25]. It has also been shown that during germination, sorghum seeds produce NO [26]. Moreover, treatment of SNP (NO source) can escalate seed germination and breaks seed dormancy [27–30]. Moreover, NO treatment promotes seed germination of wheat under elevated Cu levels by improving antioxidant capacity [31]. Keeping in mind, the reported research was undertaken to study how the exogenous application of NO could affect *B. juncea* seeds germination and antioxidant metabolism and to find out to what extent photosynthetic efficiency of Cu-treated plants was enhanced. The effect of NO was also evaluated for vegetative phase of the plant.

## Materials and methods

### Treatments

Healthy and uniform seeds of mustard (*B. juncea* L. cv. PusaTarak) were surface-sterilized with 0.1% hypochlorite (v/v) solution, washed several times with autoclaved double distilled water and the sterilized seeds were randomly placed on two filter paper covered Petri plates for germination at 28 ±2°C with 0, 0.5, 1.0, 2.0, 3.0, 4.0, 5.0, 6.0, 7.0, 8.0, 9.0 and 10.0 mM copper sulphate (CuSO_4_) for three days. Fifty seeds were placed on each Petri plate (n=50). Standard radicle emergence of seeds was scored as the germination percentage. Thus, at 3.0 mM Cu concentration the germination percentage was found declined near to half of the control and this particular concentration was considered as semi-lethal.

To study the possible and defensive role of NO donor (SNP) on seed germination under Cu stress, seeds of *B. juncea* were pre-treated with 0, 25, 50, 100, 200 and 250 μM SNP for 3 hrs, and later exposed to semi-lethal 3.0 mM Cu stress for three days. The selection of SNP concentration was done to find out its optimal concentration to alleviate Cu induced toxicity more efficiently. It was found that 100 μM SNP was optimal and most effective under 3.0 mM Cu.

In order to investigate the NO-mediated enhancement of germination of seeds under semi-lethal Cu induced stress, pretreatment of seeds to water, 100 μM SNP (optimum concentration), 100 μM potassium ferricyanide (FiCy) or 100 μM SNP plus 200 μM cPTIO (SNP + cPTIO) for 3 hrs, and later exposed to the 3.0 mM semi-lethal Cu. In this study, SNP was used as NO donor, cPTIO as NO scavenger and FiCy as SNP analogue which does not release NO.

The final experiment was conducted to evaluate whether the pretreatment of NO is limited to induce germination or it is also effective during vegetative growth of plants. To answer the question healthy and consistent seeds of mustard (*B. juncea* L. cv. Pusa Tarak) were washed and surface sterilized with 0.1% hypochlorite (v/v) and were germinated in pots filled with acid-washed autoclaved sand. The plants were raised in modified Hoagland nutrient solution applied on alternate days consisting of 3 mM KNO_3_, 0.5 μM CuSO_4_, 2 mM Ca(NO_3_)_2_, NH_4_H_3_ PO_4_, 25 μM H_3_BO_4_, 25 μM H_3_BO_4_, 2 μM MnCl_2_, 50 μM KCl, 20 μM ZnSO_4_, 20 μM Na_2_Fe-EDTA and 0.5 μM (NH_4_)_6_Mo_7_O_24_. The pots were placed in a naturally illuminated net house of the Department of Botany, Aligarh Muslim University, Aligarh (India), with an average temperature of 22 ±3°C during day and 14±2 °C at night. The treatments were arranged in a randomly blocked design with three replicates (n=3) and for each treatment. Plants were sampled at 30 days after germination (DAG) for recording different parameters. In this experiment Cu (3.0 mM) in the form of CuSO_4_ was given to seeds at the time of sowing. Seeds in the experiment were soaked or pre-treated before sowing for three hours in deionized water, 100 SNP μM, 100 μM FiCy, and 100 μM SNP plus 200 μM cPTIO. The experimental design is given in the Table 1.

**Table 1.** Experimental design

| Treatments |  |
| --- | --- |
| **Control** | Seeds pre-treated with water and raised with out Cu stress |
| **Cu** | Seeds pre-treated with water and raised with Cu stress |
| **SNP** | Seeds pre-treated with SNP and raised without Cu stress |
| **SNP + Cu** | Seeds pre-treated with SNP and raised with Cu stress |
| **FiCy + Cu** | Seeds pre-treated with FiCy and raised with Cu stress |
| **SNP + cPTIO + Cu** | Seeds pre-treated with SNP+cPTIO and raised with Cu stress |

### Determination of H_2_O_2_ content and lipid peroxidation

The assay of H_2_O_2_ was done following the method of Okuda et al. [32]. Germinated seeds (0.5 g) and fresh leaves (0.5g) of the two experiments were ground in ice-cold 200 mM perchloric acid. After centrifugation at 1,200 × g for 10 min, perchloric acid of the supernatant was neutralized with 4M KOH. The insoluble potassium perchlorate was eliminated by centrifugation at 500 × g for 3 min. The reaction was started by the addition of peroxidase and an increase in the absorbance will be recorded at A_590_ for 3 min.

Contents of TBARS were measured according to Dhindsa et al. [33] by recording absorbance at 532 nm and corrected for non-specific turbidity by subtracting the absorbance at 600 nm. The TBARS content was calculated using its extinction coefficient of 155 mM^-1^ cm^-1^.

### Assay of antioxidant enzymes

Germinated seeds (0.5 g) and fresh leaves (0.2 g) of the two experiment were homogenized with mortar and pestle by using 100 mM potassium phosphate buffer (pH 7.0) containing 1% polyvinylpyrrolidone (PVP) (w/v) and 0.05% Triton X-100 (v/v). Later the centrifugation of homogenized material was done at 15,000 × g for 20 min. The supernatant generated after centrifugation was utilized to assay the activity of glutathione reductase (GR) and superoxide dismutase (SOD). The assay of ascorbate peroxidase (APX) required addition of extraction buffer supplemented with 2 mM ascorbate.

Activity of SOD was determined as per the protocol of Beyer and Fridovich. [34] and Giannopolitis and Ries [35] by monitoring the inhibition of photochemical reduction of nitro blue tetrazolium (NBT). A 5 mL of reaction mixture consisted of 50 mM Na_2_CO_3_ (pH 10.0), 5 mM HEPES (pH 7.6), 0.1 mM EDTA, 0.025% (v/v) Triton X-100, 13 mM methionine, 63 mM NBT and 1.3 mM of riboflavin. The extract of enzyme was illuminated for 15 min (360 μmol m^2^ s^-1^), and a set was not illuminated which acted as a control to correct the turbidity of background absorbance. A unit of enzyme was expressed as the amount of the enzyme that inhibited the NBT reduction by 50% at 560 nm. The amount of the enzyme that inhibited the reduction of NBT by 50 % at 560 nm was equal to one unit of SOD.

Activity of APX and GR was assessed as per the protocol adopted by Nakano and Asada. [36] and Foyer and Halliwell. [37]. The detailed procedure for determining APX and GR activity has been explained in our earlier study [38].

### Assay of amylase activity

Seeds were pre-treated with water (control) or with SNP (100 μM), later pre-treated seeds were kept for germination for 48 hrs in Petri plates containing 3.0 mM CuSO_4_ solution. Seeds (0.5 g) were then collected and homogenized in a pre-chilled mortar and pestle with 6 mL of 50 mM Tris–HCl (pH 7.5) containing 1% PVP and 15 mM 2-mercaptoethanol. The homogenate was collected and centrifuged at 4°C at 10,000 × g for 30 min. After centrifugation, the resulting supernatant was used for determining enzyme activity. The amylase activity was determined by using the protocol of starch-iodine following Collins et al. [39]. A unit of enzyme was expressed by taking the enzyme quantity to reach 50% of the original color intensity.

### Determination of reduced glutathione (GSH) content

GSH content in fresh leaves was determined spectrophotometrically according to the protocol provided by Anderson. [40]. Using pre-chilled mortar and pestle, under 4°C fresh leaf tissue (0.5 g) was crushed in 2.0 mL of 5% sulphosalicylic acid. Centrifugation of homogenized material was done at 10,000 × g for 10 min. Then, 0.6 mL of phosphate buffer (100 mM, pH7.0) and 40 mL of DTNB were added to the 0.5 mL of supernatant. After two minutes, the absorbance was recorded at 412 nm. The detailed protocol has been given in our earlier publication [41]

### Histochemical detection of ROS

Histochemical staining was performed using the protocol given by Kumar et al. [42]. Nitro blue tetrazolium (NBT) and 3, 3-Diaminobenzidine (DAB) were used for the assay accumulation of both superoxide ion (O_2_^.-^) and H_2_O_2_ in seed and leaf samples of different treatments. The samples from each treatment were kept into NBT solution prepared by dissolving 0.1 g of NBT in 50 mL of 50 mM sodium phosphate buffer (pH 7.5) in an amber-colored bottle and were incubated overnight. The stained samples were immersed in absolute ethanol and boiled in water-bath for 10 min for discoloration to get the staining clear.

For DAB staining solution, 50 mg DAB was dissolved in 50 ml double-distilled water in an amber-colored bottle with pH 3.8. The samples from each treatment were kept into DAB solution and incubated it for 8 hrs. The stained samples were immersed in absolute ethanol and boiled in a water bath for 10 min for discoloration to get a clear visualization of the stained samples.

### Confocal laser microscopy study for ROS imaging and cell viability determination

For ROS imaging, root samples 1-2 cm in length were dipped into the 12.5 μM 2′,7 ′ Dichlorofluorescindiacetate (H_2_DCFDA) solution in a petri-dish for 15 min and washed three times properly with double distilled water. Stained samples were kept on a glass slide and studied under a Confocal Microscope (excitation 400–490nm, emission ≥ 520nm).

For the cell viability test, properly washed root samples (1-2 cm length) of each treatment were dipped into 25 μM propidium iodide (PI) solution. Stained samples were washed appropriately and placed on a glass slide and were observed using a confocal microscope.

### Determination of Cu concentration

Leaf and root samples were dried in the oven for two days at 80°C. The oven-dried samples were crushed to a fine powder in mortar and pestle. This fine powder was digested with a solution containing concentrated HNO_3_/HClO_4_ (3:1, v/v) and was diluted with water. Concentration of Cu was determined by Atomic Absorption Spectrophotometer (GBC, 932 plus; GBC Scientific Instruments, Braeside, Australia).

### Photosynthetic characteristics

Net photosynthesis (P_N_), intercellular CO_2_ (C_i_) and stomatal conductance (g_S_) were measured in second topmost leaves of plants using Infrared gas analyzer (CID-340, Photosynthesis System, Bio-Science, USA). These parameters were recorded between 11 am to 12 noon when the photosynthetically active radiation (PAR) was above 780 μmol m^-2^ s^-1^ and at 380 ± 5 μmol^-1^ atmospheric CO_2_ concentrations.

Maximal quantum efficiency of photosystem II (PSII) (*Fv/Fm*) of full-fledged leaf second from the top was noted with the help of chlorophyll fluorometer (Heinz Walz, Germany). Preceding to get the result of maximum fluorescence (*Fm*) and minimal fluorescence (*Fo*) intensity, leaf samples were kept in the dark condition for 30 minutes. Saturating pulse (720 µmol m^-2^ s^-1^) and weak measuring pulses (125 µmol m^-2^s^-1^) were used to measure *Fo* and *Fm* respectively. Difference between *Fo* and *Fm* was used to calculate the variable fluorescence (Fv). The maximum quantum yield efficiency of PS II was calculated as a ratio of *F_v_ t*o *F_m_*.

Activity of Rubisco in leaves was monitored by adopting the procedure of Usuda, [43]. One gram of fresh leaf samples were ground in a pre-chilled mortar and pestle with an extraction buffer containing 0.25 M Tris-HCl (pH 7.8), 0.0025 M EDTA, 0.05 M MgCl_2_ and 37.5 mg dithiothreitol (DTT). Centrifugation of homogenized material was done at 10,000 × g for 10 min at 4 °C. The resulting supernatant brought after centrifugation was used to obtain measure enzyme activity. A reaction mixture (3 ml) was formed by mixing 100 mM Tris-HCl (pH 8.0), 10 mM MgCl_2_, 0.2 mM NADH, 40 mM NaHCO_3_, 5 mM DTT, 4 mM ATP, 1U of 3-phosphoglycerate kinase, 1U of glyceraldehyde 3-phosphodehydrogenase, and 0.2 mM ribulose 1, 5-bisphosphate (RuBP). Bradford [44] method was adopted to estimate protein content.

### Determination of growth parameters

The plant samples from each treatment were dried in a hot air oven at 80°C. The dried leaf samples were weighed on an electrical balance and the weight was recorded as whole plant dry mass. To measure leaf area, leaf area meter (LA 211, Systronics, New Delhi, India) was used.

### Physiological measurements of guard cells

Upper leaves of 30-day-old plants were plugged from each plant with different treatments and were fixed by 2.5% glutaraldehyde, and stomatal images were taken using scanning electron microscopy (JSM-6510 LV, JEOL, Japan)

The epidermal peels of the leaf samples were removed from the abaxial side and stomatal images of the sections at 40x were taken using a compound microscope outfitted with NIKON digital camera. The stomatal aperture width was measured with the help of a micrometer scale.

### NO generation

The level of NO generation was confirmed by estimating nitrite content adopting the protocol given by Zhou et al. [45] with slight modifications. Using pre-chilled mortar and pestle, leaf samples (500 mg) were ground in 3 ml of 50 mM ice-cold acetic acid buffer (pH 3.6) containing 4% zinc acetate. Later, centrifuged at 11,500 × g for 15 min at 4°C. The pellet obtained was washed twice with 1 ml of the extraction buffer and then centrifuged again. The resulting supernatants from the two spin were mixed and neutralized by the addition of 100 mg of charcoal. After brief vortex, the filtrate was collected. Each filtrate of 1 ml and Greiss reagent (0.1% N-1-naphthyl ethylenediamine dihydrochloride and 1% sulphanilamide in 5% H_2_PO_4_ solution) were mixed in the ratio of (1:1) and then for 30 min these were incubated at room temperature. The absorbance was read at 540 nm and NO content was measured from a calibration curve plotted using sodium nitrite as standard.

### Statistical analysis

Data in the experiments were analyzed statistically using analysis of variance (ANOVA) by SPSS 17.0 for Windows, and presented as treatment mean ± SE (n = 3). Treatment means were compared using the least significant diifernce (LSD) at P < 0.05. Bars showing the same letter are not significantly different by LSD test at P < 0.05.

## Results

### Effect of Cu stress on seed germination

The germination percentage of seeds declined with the increasing Cu concentration of from 1 to 10 mM (Fig 1A). At 3.0 mM Cu concentration, germination percentage declined near to half of the control and was marked as a semi-lethal concentration and later this Cu concentration was selected for further experimentations. Fig 1(B) offers an overview showing Cu induced inhibitory effect on seed germination under various Cu concentrations.

**Fig 1.**
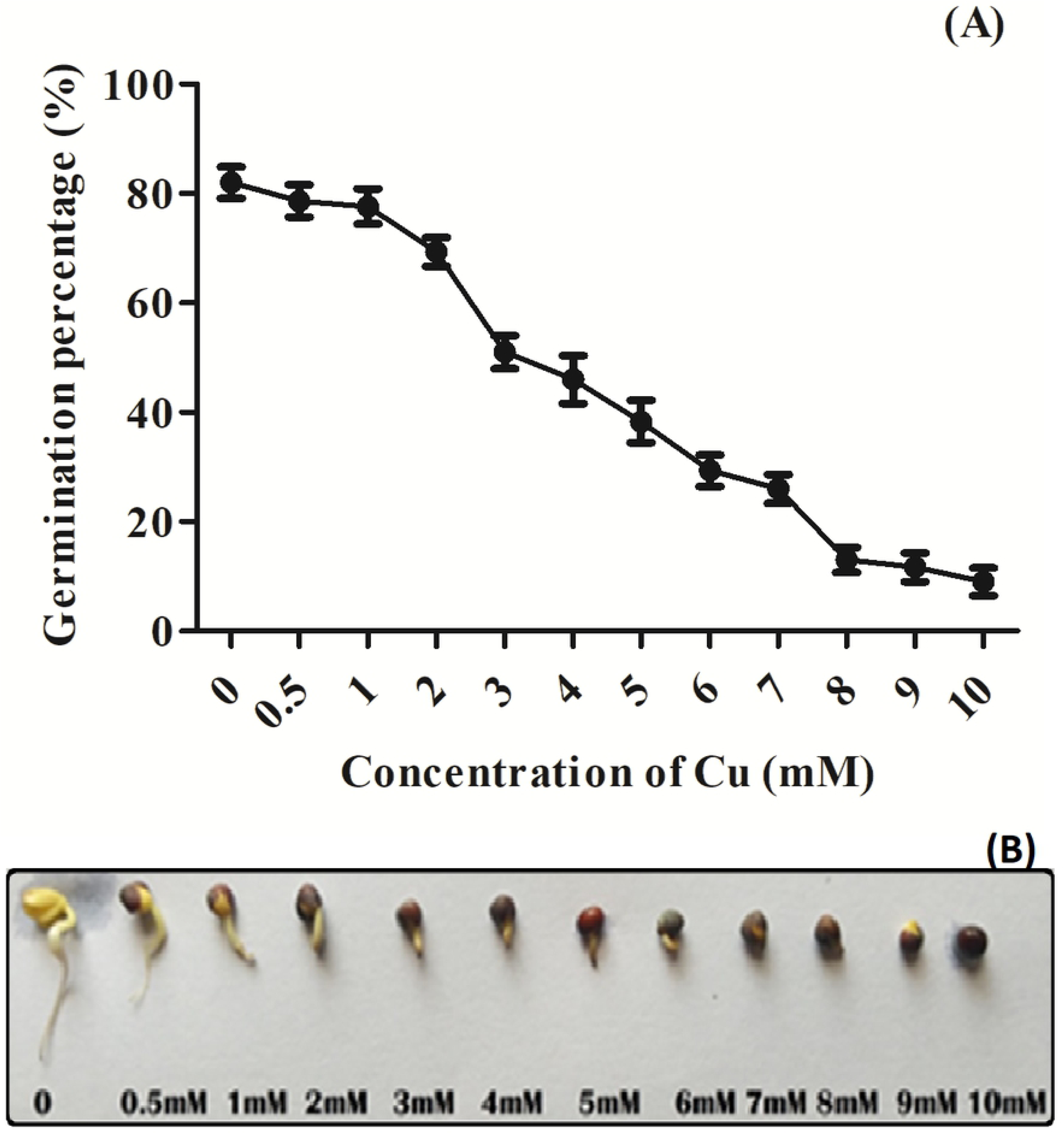
**(A)** Germination percentage; **(B)** morphology of randomly selected germinating mustard seeds; Germination percentage in mustard seeds (*B. juncea* L.) exposed to copper (Cu); (0, 0.5, 1.0, 2.0, 3.0, 4.0 5.0, 6.0, 7.0, 8.0, 9.0 and 10.0 mM) for 3 days.

### Effect of NO on seed germination under Cu stress

Seeds pre-treated with 0, 25, 50, 100, 200, and 250 µM SNP concentration for 3 hrs were later exposed to 3.0 mM Cu showed increased germination percentage up to 100 µM SNP, then dropped to approximately the same as control at 250 µM. Here, 100 µM SNP was proved to be the most effective treatment against 3.0 mM Cu stress-induced inhibition of seed germination (Fig 2A). Fig 2(B) is a critique of seed germination under varying SNP concentrations.

**Fig 2.**
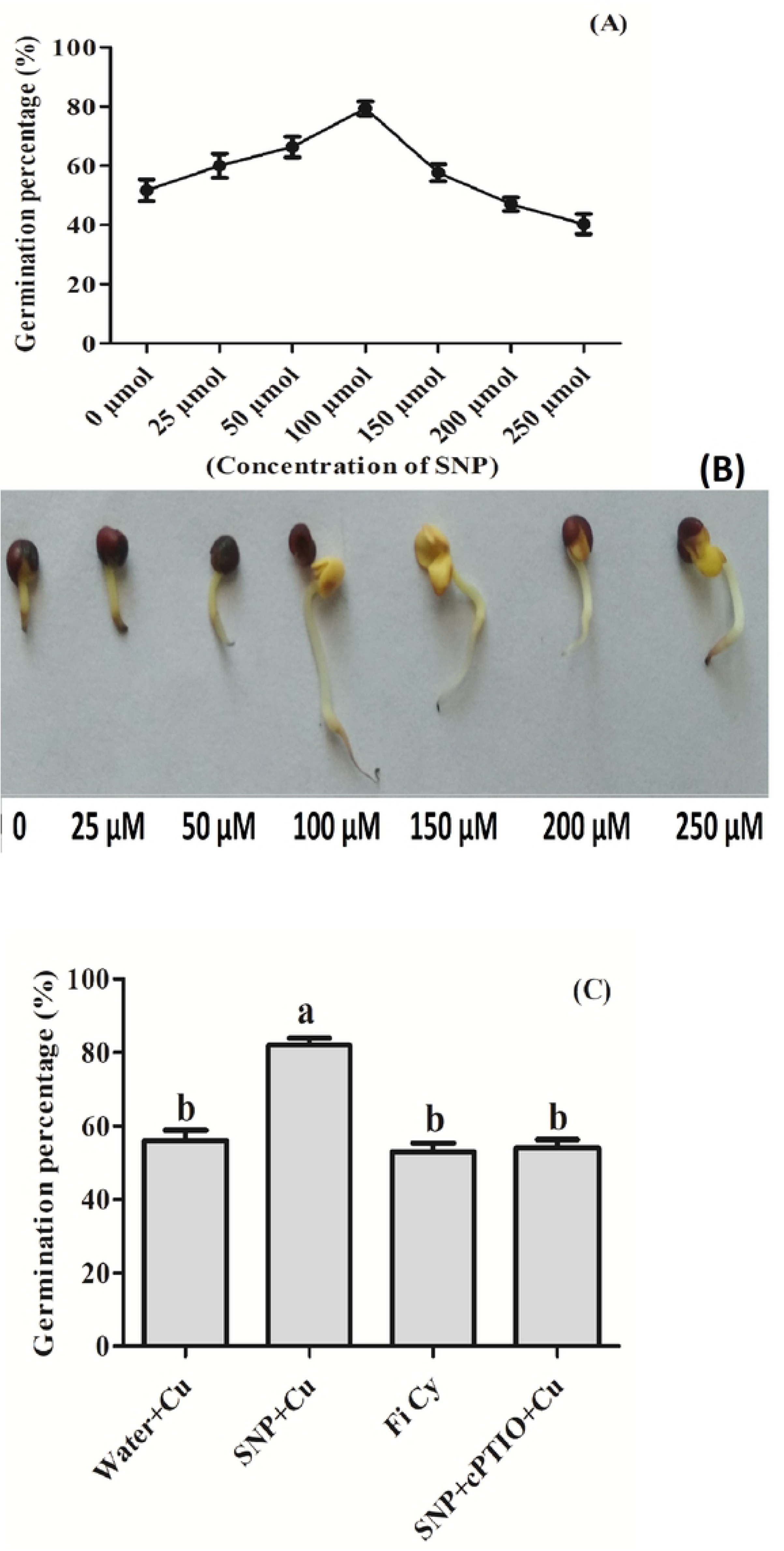
**(A)** Germination percentage and **(B)** morphology of randomly selected germinating seeds of mustard (*B. juncea* L.) Pre-treated with 0, 25, 50, 100,150.200 and 250 µM SNP and exposed 3.0 mM Cu; and (**C)** germination percentage in Mustard seeds (*B. juncea* L,) pretreated with water, SNP, SNP + cPTIO (cPTIO as a specific NO scavenger), and FiCy (SNP analouge) exposed to 3.0 mM Cu stress for 3 days.

SNP pretreatment showed an increase in seed germination percentage under Cu stress. But the pretreatment of Cu exposed seeds to NO analogue FiCy showed no significant change in germination percentage in comparison to Cu exposed seeds. Further, the use of cPTIO along with SNP in Cu exposed seeds reversed the SNP induced inhibition of Cu stress effect on seed germination. From these results, it can be assumed that the NO released by SNP was behind the reversal of seed germination inhibition in seeds of mustard plants, since not only its analogue worked in mitigation but also inhibition of NO generation using cPTIO was proved ineffective (Fig 2C).

### Effect of NO on H_2_O_2_ and TBARS contents in germinating seeds under Cu stress

Application of SNP to seeds prior germination under Cu stress reduced H_2_O_2_ content by about 61.2% and TBARS 51.6% in comparison to the water pretreated germinating control Cu stressed seeds. However, FiCy and SNP in combination with cPTIO pretreatment reduced 9.8% and 9.7% H_2_O_2_ content, respectively as compared to control (Cu stressed only). In case of TBARS content FiCy and SNP in combination with cPTIO reduced it by 9.3% and 3.8% respectively in comparison to water pretreated control seeds (Fig 3A and B).

**Fig 3.**
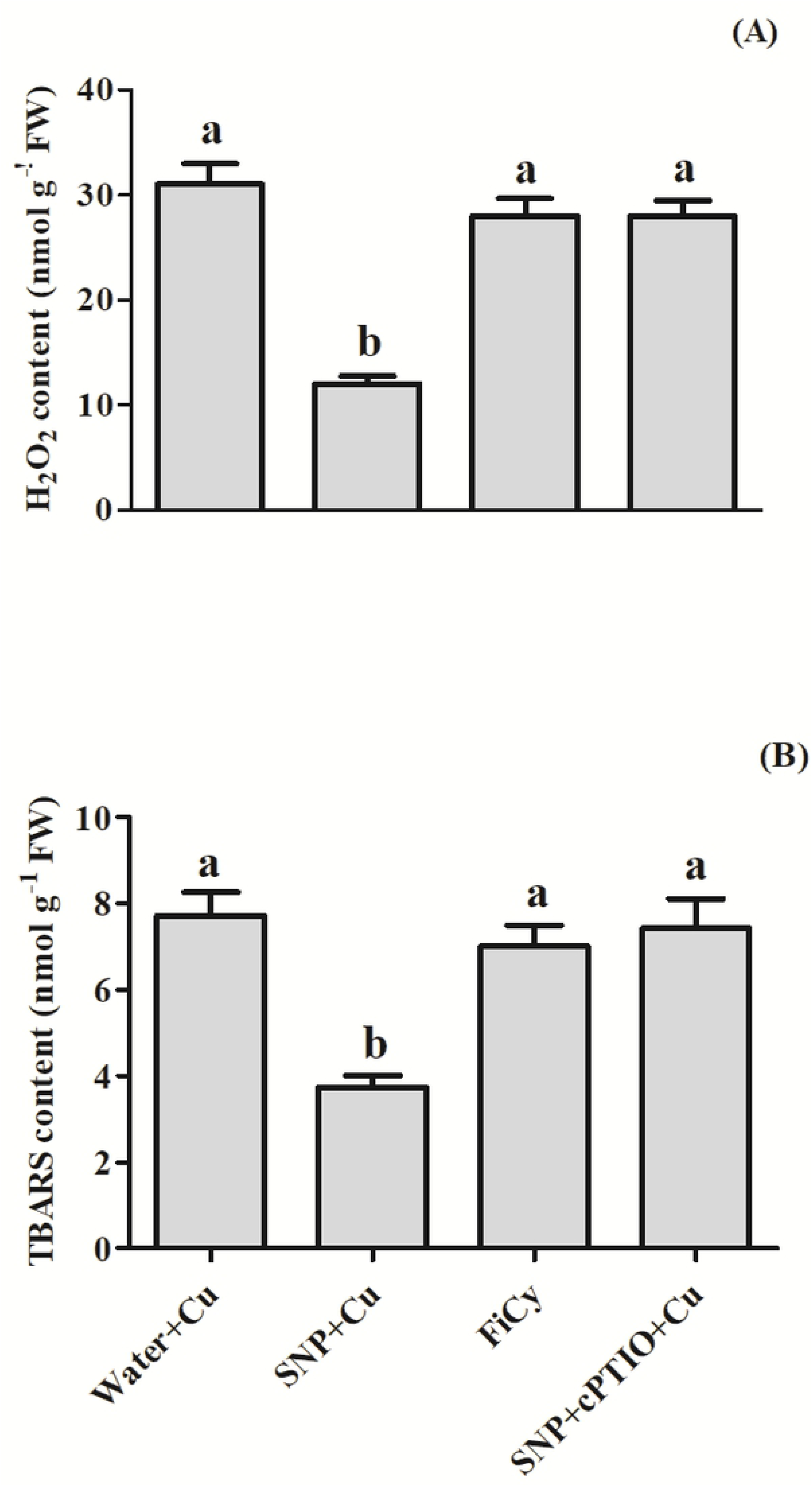
H_2_O_2_ (**A**) and TBARS (**B**) content in germinating seeds of mustard (*B. juncea* L.) pre-treated with water, SNP, SNP + cPTIO (cPTIO as a specific NO scavenger), and FiCy (SNP analouge) exposed to 3.0 mM Cu stress for 3 days. Same letter above bars show that data did not differ significantly by LSD test at P < 0.05.

### Effect of NO application on activities antioxidant enzymes under Cu stress during seed germination

SNP in the presence of Cu boosted the activity of antioxidant enzymes to a higher level than water pretreated germinated control seeds; increasing the activity of APX enzyme by about 111.7%, SOD enzyme by about 81.8 %, and GR by 233.9% in contrast to the control seeds. However, in the presence of Cu, SNP in combination with cPTIO showed no difference than the control in the activity of antioxidant enzymes. Similarly, FiCy also showed significantly similar activity of APX, SOD and GR as was under Cu treatment alone (Fig 4).

**Fig 4.**
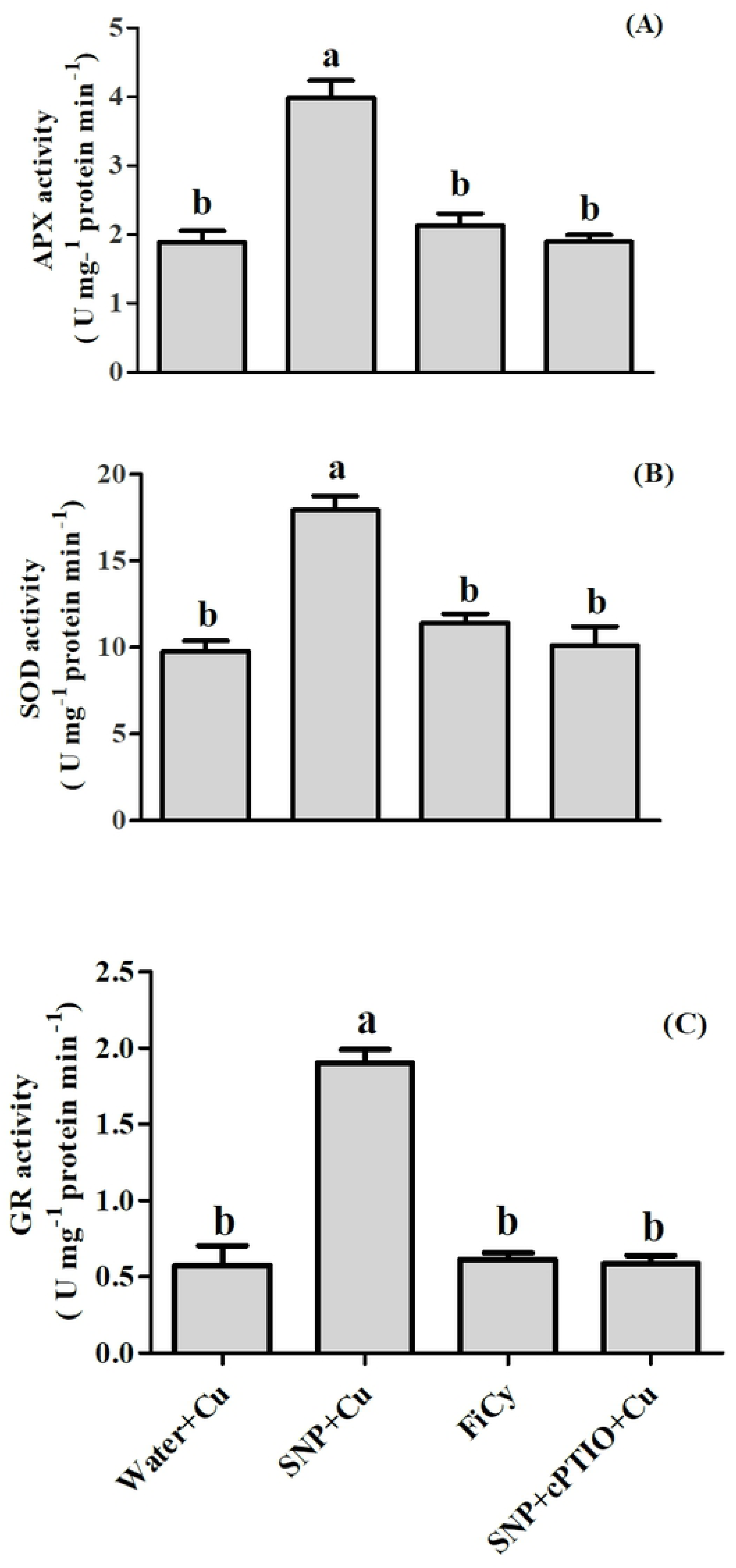
Activity of APX (**A**), SOD (**B**) and GR (**C)** in germinating seeds of mustard (*B.juncea* L.) pre-treated with water, SNP, SNP + cPTIO (cPTIO as a specific NO scavenger), and FiCy (SNP analouge) exposed to 3.0 mM Cu stress for 3 days.

### ROS accumulation in germinating seeds

In seeds pretreated with water, FiCy and SNP in combination with cPTIO showed an increase in O_2_^.-^ accumulation as compared to SNP pretreated seeds. Which was observed by histochemical staining with NBT, showed dark blue stained area on radicle (a marker for O_2_^.-^). While in case of accumulation of H_2_O_2_ after germination which was revealed by DAB staining showed more reddish-brown precipitate on water pretreated seeds, FiCy and SNP + cPTIO pretreated seeds as compared to SNP treatment under Cu stress (Fig 5A and B).

**Fig 5.**
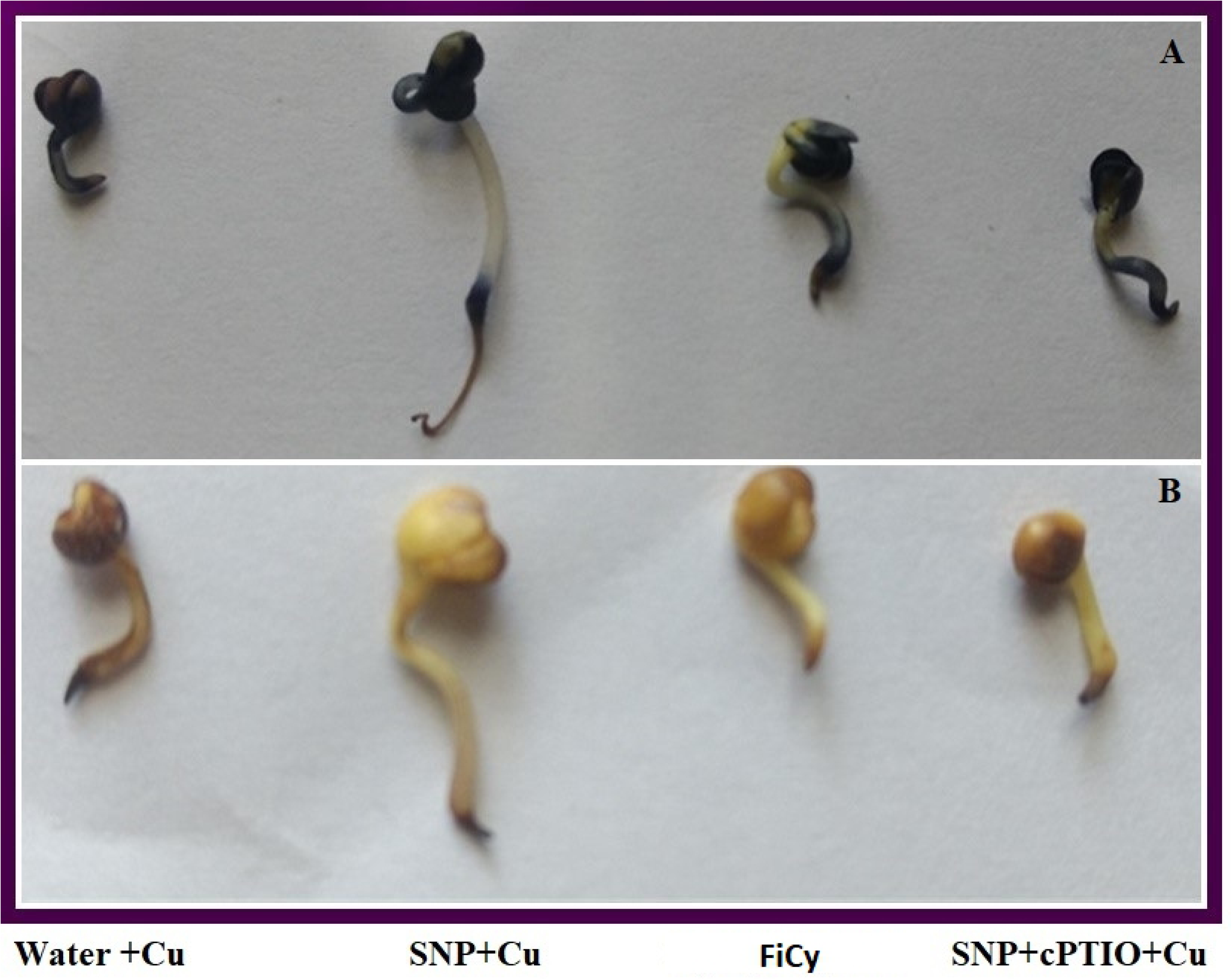
Accumulation of superoxide ion (O_2_^.-^) by NBT (**A**) and H_2_O_2_ by DAB staining (**B**) of germinating mustard seeds (*B. juncea* L.) pre-treated with water, SNP, SNP + cPTIO (cPTIO as a specific NO scavenger), and FiCy (SNP analouge) exposed to 3.0 mM Cu stress for 3 days.

### Effect of NO on the activity of amylase

From Fig 6, it can be observed that during 48 hrs of germination, activity of amylase changes with the duration of treatment. SNP pretreated seeds have higher amylase activity as compared to the control with a continuous rise to a maximum till 24 hrs under Cu stress. Before 24 hrs amylase activity increased in both the treatments, but after 24 hrs amylase activity gradually declined.

**Fig 6.**
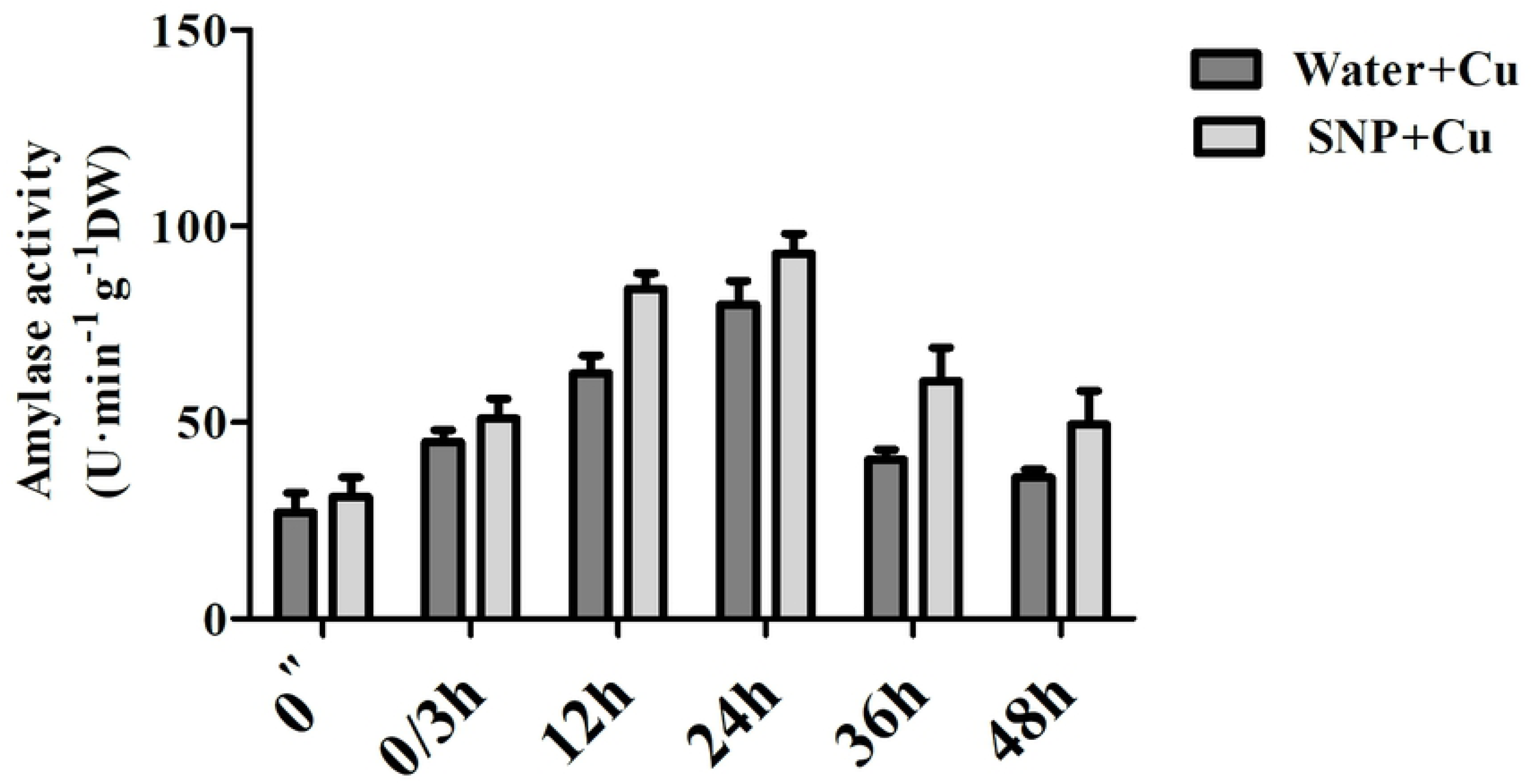
Activity of amylase in germinating seeds of mustard (*B. juncea* L.) pre-treated with water as control or 100 µM SNP for 3 h (shown as from 0” to 0/3h of the treatment times) prior to exposing to 3.0 mM Cu stress for further 48 h (shown as 3/0, 12, 24, 36, 48 h respectively).

### Effect of NO on Cu uptake and levels of H_2_O_2_ and TBARS

To examine the possible role of NO in the *B. juncea* plant response to Cu toxicity, we measured the Cu content in roots and leaves in Cu exposed plants (Fig 7A, B). The Cu content accumulation was much higher in roots (470 µg g^-1^) than in leaves (28.02 µg g^-1^). Pre-germination application of NO in the form of SNP resulted in a decrease in Cu content level in both roots and leaves of Cu treated plants. However, the protecting effect of NO against Cu toxicity was reversed by NO scavenger, cPTIO when applied in combination with SNP.

**Fig 7.**
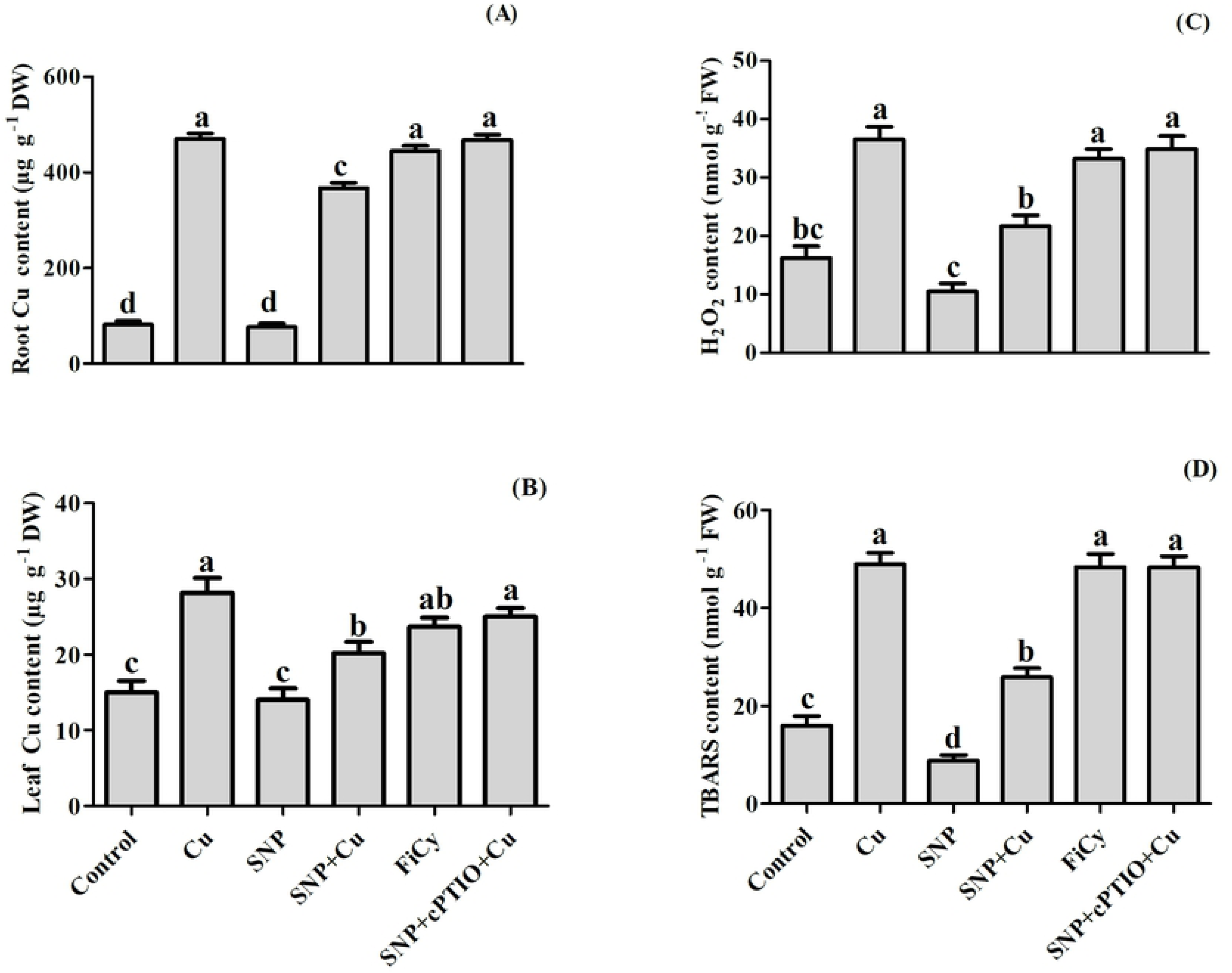
Contents of Cu in root (**A**) and leaf (**B**), and contents of H_2_O_2_ (**C**) and TBARS (**D**) in mustard (*B. juncea* L.) treated during pre-germination with water, 100 µM SNP or SNP with cPTIO, FiCy in the presence or absence of 3 mM Cu at 30 DAG. Same letter above bars show that data did not differ significantly by LSD test at P < 0.05.

Compared to control, plants receiving Cu showed greater content of H_2_O_2_ and TBARS (Fig 7 C, D). The individual application of NO in the form of SNP exhibited decrease in H_2_O_2_ and TBARS content by about 35% and 45% without Cu stress, but this decrease due to NO in H_2_O_2_ and TBARS content was by about 40%, and 47% in Cu stressed plants in comparison with Cu-treated plants. The protective effect of NO application was blocked by addition of cPTIO in Cu fed plants. Further, FiCy effect was significantly similar to the effect generated on application of cPTIO to SNP and Cu treated plants.

### Effect of NO on ROS accumulation

In our experiment, both H_2_O_2_ and O_2_^-^ markedly accumulated in Cu stressed-plant (Fig 8 A-F and G-L). Accumulation of H_2_O_2_ and O_2_^-^ was shown by histochemical staining with DAB and NBT, respectively. The leaves from the Cu treated plants, with FiCy and cPTIO in combination with NO applied plants showed discrete and deepest blue staining which is a marker of O_2_^-^ accumulation. However, adding SNP to non-stressed seeds seemed closer to control, although application of NO noticeably diminished ROS accumulation in the leaves of Cu-stressed *B. juncea* leaves. A similar result was observed in DAB staining, but in this case, brownish patches were observed which was a marker of H_2_O_2_ accumulation in leaves.

**Fig 8.**
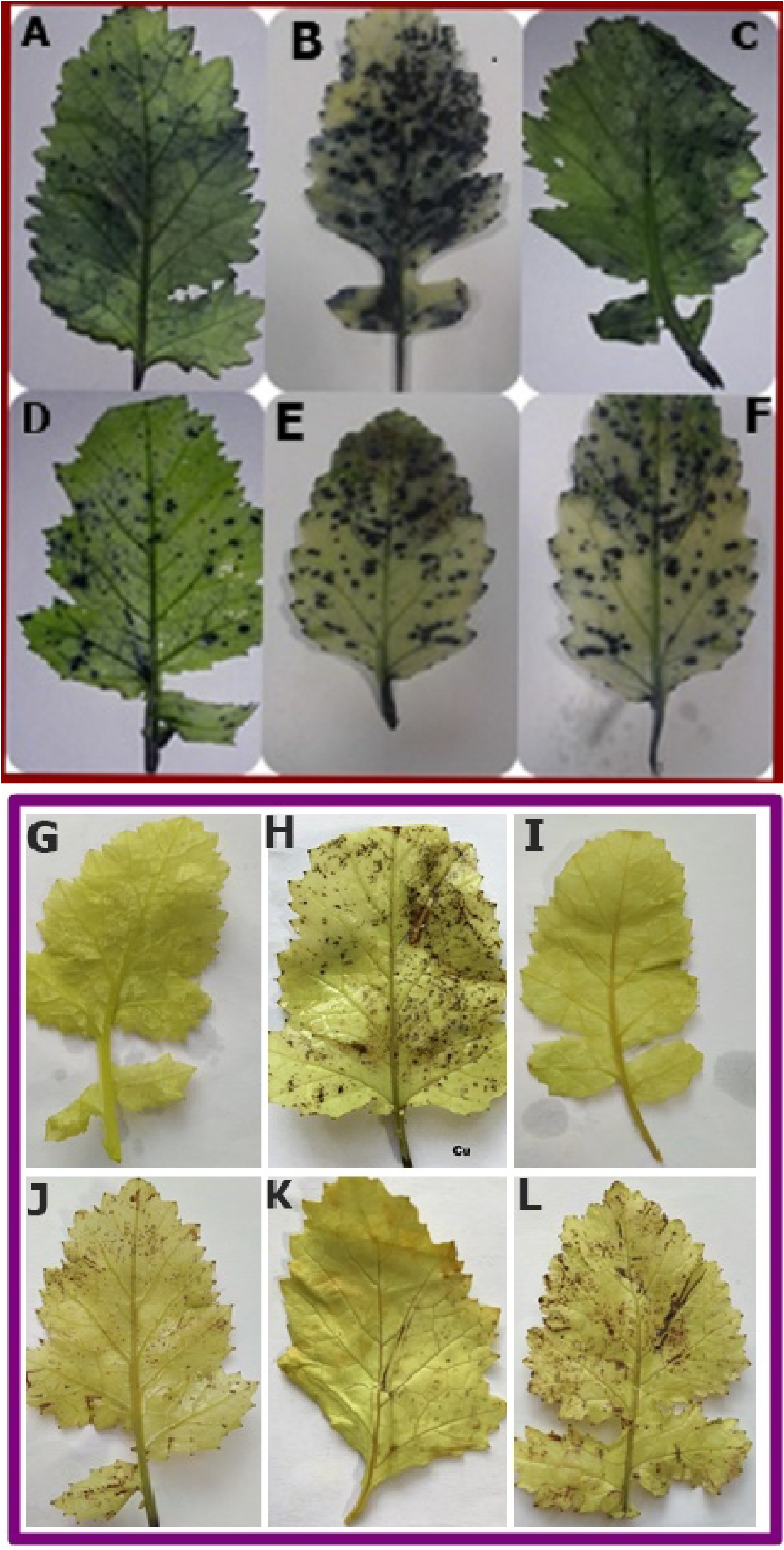
Accumulation of superoxide ion (O_2_^.-^) and H_2_O_2_ in leaves stained with by NBT (**A-F**) and DAB (**G-L**) respectively of mustard (*B. juncea* L.) treated during pre-germination with water, 100 µM SNP or SNP with cPTIO, FiCy in the presence or absence of 3 mM Cu at 30 DAG. Control: **A** and **G**; **B** and **H**: Cu; **C** and **I**: SNP; **D** and **J**: SNP+Cu; **E** and K: FiCy; **F** and **L**: SNP+cPTIO+Cu

### Confocal laser scanning microscopy

Cu induced H_2_O_2_ generation was visualized in roots by staining with H_2_DCFDA an indicator for ROS predominantly H_2_O_2_ in cells (Fig 9 A-F). This reagent passively diffused into cells and its acetate groups cleaved by esterases. Upon oxidation, by H_2_O_2_ the non-fluorescent probe H_2_DCFDA gets converted to the highly fluorescent 2’, 7-dichlorofluorescein (DCF). In our study root cells of Cu stressed and FiCy or SNP in combination with cPTIO treated plants in the presence of Cu yielded higher intensity of green fluorescence while SNP reduced the effect of Cu stress and showed the less intensity of green fluorescence like control plant (Fig 9. A-F)

PI staining (an indicator of cell death) used to visualize cell viability by identifying the nucleic acid staining. PI is membrane impermeant and generally excluded from viable cells and in dead cells reaches the nucleus through distorted areas of dead cell membranes. In our study root cells of Cu stressed, FiCy and of SNP in combination with cPTIO were less viable. However, Cu induced cell death was reduced by SNP application, showing similar response to control plants (Fig 9. G-L).

**Fig 9.**
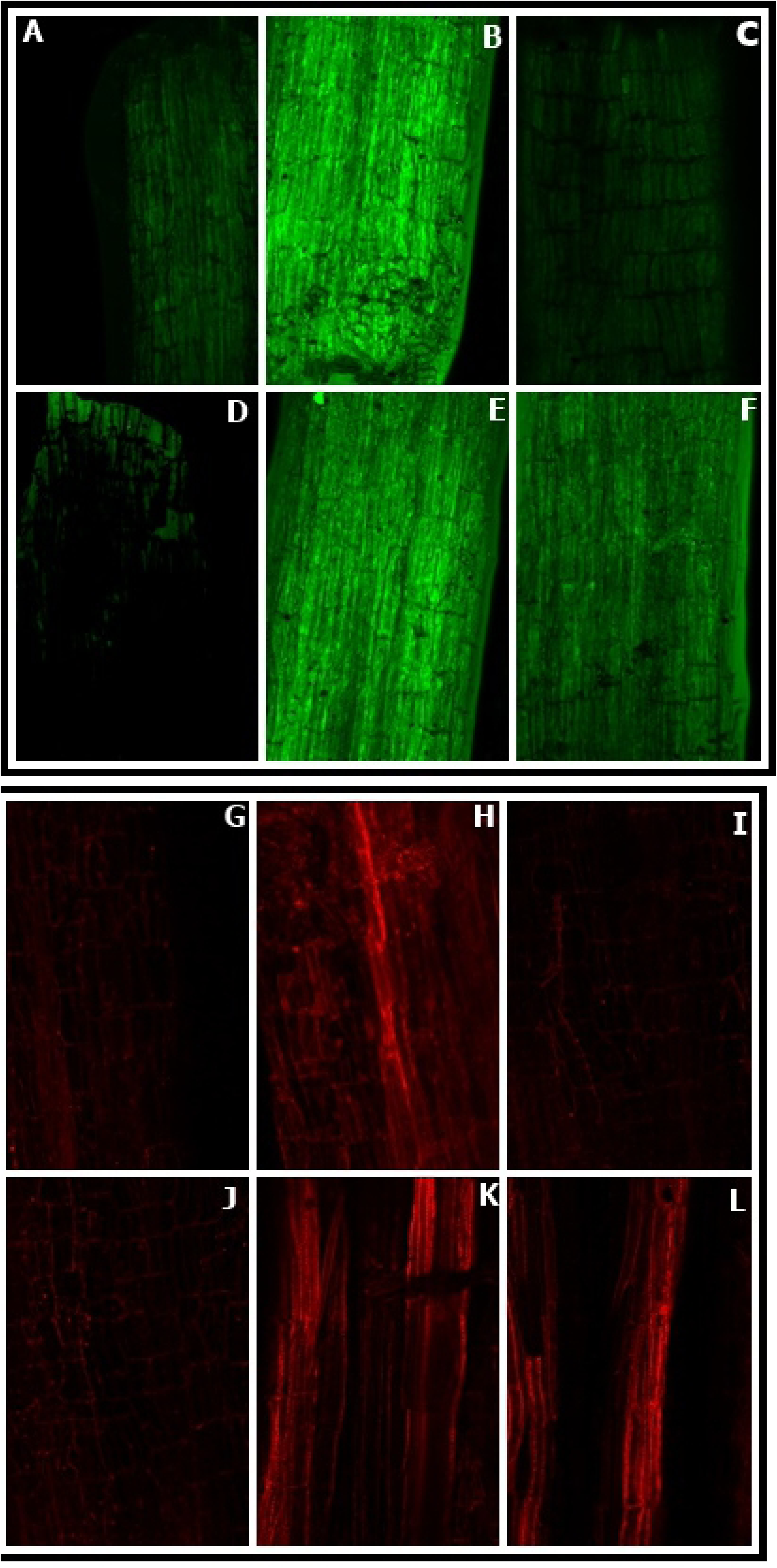
Confocal microscopic images of H_2_O_2_ formation in roots using H_2_DCFDA staining (**A-F**) and Cell viability test (**G-L**) by Propidium iodide (PI) staining was performed on 30 days old roots of mustard (*B. juncea* L.) treated during pre-germination with water, 100 µM SNP or SNP with cPTIO, FiCy in the presence or absence of 3 mM Cu at 30 DAG. Control: A and G; **B** and **H**: Cu; **C** and **I**: SNP; **D** and **J**: SNP+Cu; **E** and **K**: FiCy; **F** and **L**: SNP+cPTIO+Cu.

### Effect of NO on the activity of antioxidant enzymes

In Fig 10, it is apparent that in leaves of *B. juncea* SOD activity increased with Cu treatment in contrast to the control. SNP enhanced the SOD activity to a higher level than in the presence of Cu alone. However, pretreatment of SNP in stressed plant maximally elevated SOD activity, which was by 99.6% when compared to the control. FiCy and cPTIO in combination with SNP did not alter the SOD enzyme activity in the stressed plant and showed response comparable to control.

**Fig 10.**
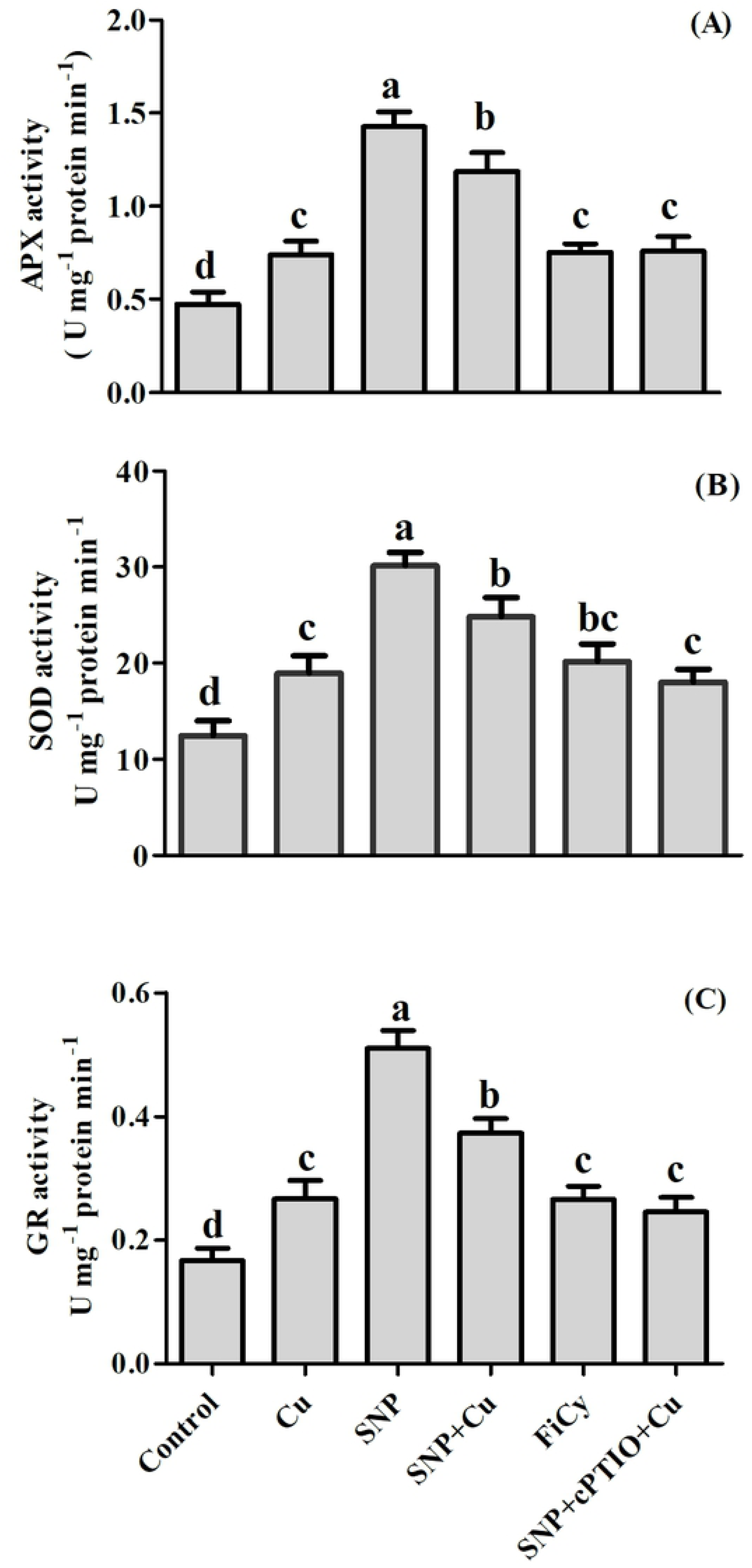
APX (**A**), SOD (**B**), and GR (**C**), activity in leaves of mustard (*B. juncea* L.) treated during pre-germination with water, 100 µM SNP or SNP with cPTIO and FiCy in presence or absence of 3 mM Cu at 30 DAG.. Same letter above bars show that data did not differ significantly by LSD test at P < 0. 05.

Cu treatment increased the activity of GR and APX by 60% and 56.3% respectively as compared to control. Maximum increase in GR and APX activity was noted with the supplementation of NO (Cu + SNP) which increased the activity of both by about 40% and 60% respectively in leaves in comparison to Cu treated plant. While by addition of NO scavenger cPTIO nullified the influence of NO. Apart from that FiCy had also a null effect on antioxidant enzymes activities in the Cu exposed plant.

### Impact of NO on photosynthetic performance

Compared to the control plants getting NO in the form of SNP exhibited higher values for photosynthetic characters such as P_N_, Ci, gs, maximal PSII photochemical efficiency, and Rubisco activity. Supplementation of NO in non-Cu fed plants improved P_N_ value by 57%, Ci by 49.3%, gs by 36.2%, maximal PSII photochemical efficiency by 33.3% and Rubisco enzyme activity by 35.0% when compared to the control. However, Cu treatment reduced the values of above parameters by 38%, 21.7%, 29.5%, 45.1%, 40.7%, respectively. Further, in Cu-treated plants, NO alleviated Cu induced reduction and increased P_N_, Ci, gs, and PSII photochemical efficiency and Rubisco enzyme activity by 114%, 74.8%, 84.4%, 114.7%, 89%, respectively in contrast to the Cu treated plants. However, in case of supplementation of FiCy the results were same for all photosynthetic parameters as shown by the Cu stressed plant. While treatment of cPTIO in combination with NO nullified the effect of NO, resulted in the removal of the effect of NO in Cu exposed plants (Table 2).

**Table 2.** Net photosynthesis, intercellular CO_2_ concentration, Maximal PSII photochemical efficiency, stomatal conductance, and Rubisco activity of Pusa Tarak cultivar of mustard (*Brassica juncea* L.) at 30 DAG. Seeds were treated at pre-germination time with water, 100 µM SNP or SNP with cPTIO and FiCy in presence or absence of 3 mM Cu.

| Treatments | Net Photosynthesis ($P_N$) ($\mu mol CO_2 m^{-2} S^{-1}$) | Internal $CO_2$ Concentration ($C_i$) ($\mu mol CO_2 mol^{-1}$) | Maximal PSII Photochemical efficiency | Stomatal conductance ($g_s$) ($mmol CO_2 m^{-2} S^{-1}$) | Rubisco Activity ($\mu mol CO_2 mg^{-1} protein min^{-1}$) |
| --- | --- | --- | --- | --- | --- |
| Control | 16.67 ± 0.89<sup>c</sup> | 203±5.57<sup>c</sup> | 0.6 ± 0.09<sup>c</sup> | 288±9.54<sup>b</sup> | 0.82 ± 0.06<sup>b</sup> |
| Cu | 10.27 ± 0.92<sup>e</sup> | 159±5.51<sup>d</sup> | 0.34 ± 0.06<sup>d</sup> | 203±8.54<sup>c</sup> | 0.49 ± 0.05<sup>c</sup> |
| SNP | 26.16 ± 0.72<sup>a</sup> | 303±4.04<sup>a</sup> | 0.8 ± 0.08<sup>a</sup> | 392.67±8.19<sup>a</sup> | 1.06 ± 0.07<sup>a</sup> |
| SNP + Cu | 22 ± 0.58<sup>b</sup> | 278±6.08<sup>b</sup> | 0.7 ± 0.06<sup>b</sup> | 374.33±8.68<sup>a</sup> | 0.92 ± 0.09<sup>ab</sup> |
| FiCy + Cu | 12.96 ± 0.80<sup>d</sup> | 168±5.29<sup>d</sup> | 0.38 ± 0.07<sup>d</sup> | 221±6.66<sup>c</sup> | 0.52 ± 0.04<sup>c</sup> |
| SNP + cPTIO + Cu | 11.97 ± .58<sup>de</sup> | 162±4.51<sup>d</sup> | 0.35 ± 0.05<sup>d</sup> | 208±5.51<sup>c</sup> | 0.51 ± 0.04<sup>c</sup> |

### Effect of NO on physiology of guard cells

Changes in stomatal guard cells in response to SNP, FiCy and cPTIO were studied under electron microscopy. Stomatal analysis depicted a noteworthy decrease in the stomatal within leaves of Cu exposed plants as compared to the control. Stomatal opening (length and width) was 7.96 and 0. 92μm on 30^th^ DAG of control plants (Fig 11), while it was about 9.42 μm in pore length and stomata was closed in leaf samples of Cu exposed plants. However, the supplementation of SNP in the presence of Cu showed an increase in the stomatal aperture by about 2.29 μm in diameter. Conversely, treatment of SNP applied plants resulted in the maximal stomatal opening of 2.53 and 12.54 μm in pore length and pore width respectively. Further, the application of SNP treated Cu-stressed plants with its scavenger cPTIO showed decrease in the stomatal aperture by about 0.81 μm in diameter. However, when FiCy was applied to Cu-treated plants showed the same result as plants grown under Cu treatment alone. (Fig 11, A-F.)

**Fig 11.**
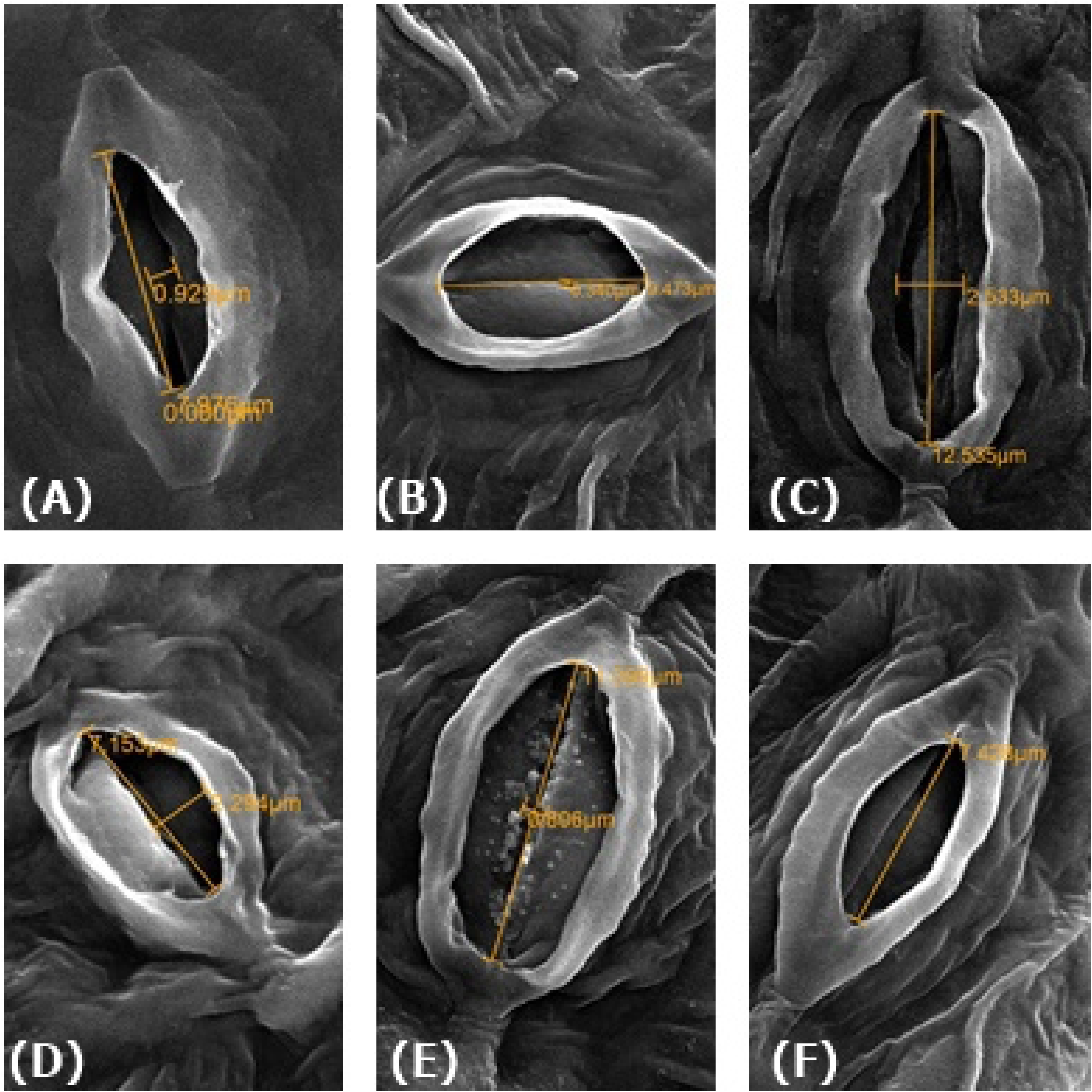
Stomatal response of *B. juncea* L. plants at 30 DAG under control **(A)**, Cu **(B)**, SNP **(C)**, SNP **+ Cu (D)**, FiCy+ Cu (**E)** and SNP **+** Cu +cPTIO **(F)** at 4000× using scanning microscope.

The assertions of electron microscopy were further expanded by compound microscopy observations. Stomata in leaf samples of control were normal with specialized guard cells, SNP applied in Cu treated plants showed an increase in stomatal aperture as compared to closed with distorted guard cells in stressed plants (Fig 12, A-F)

**Fig 12.**
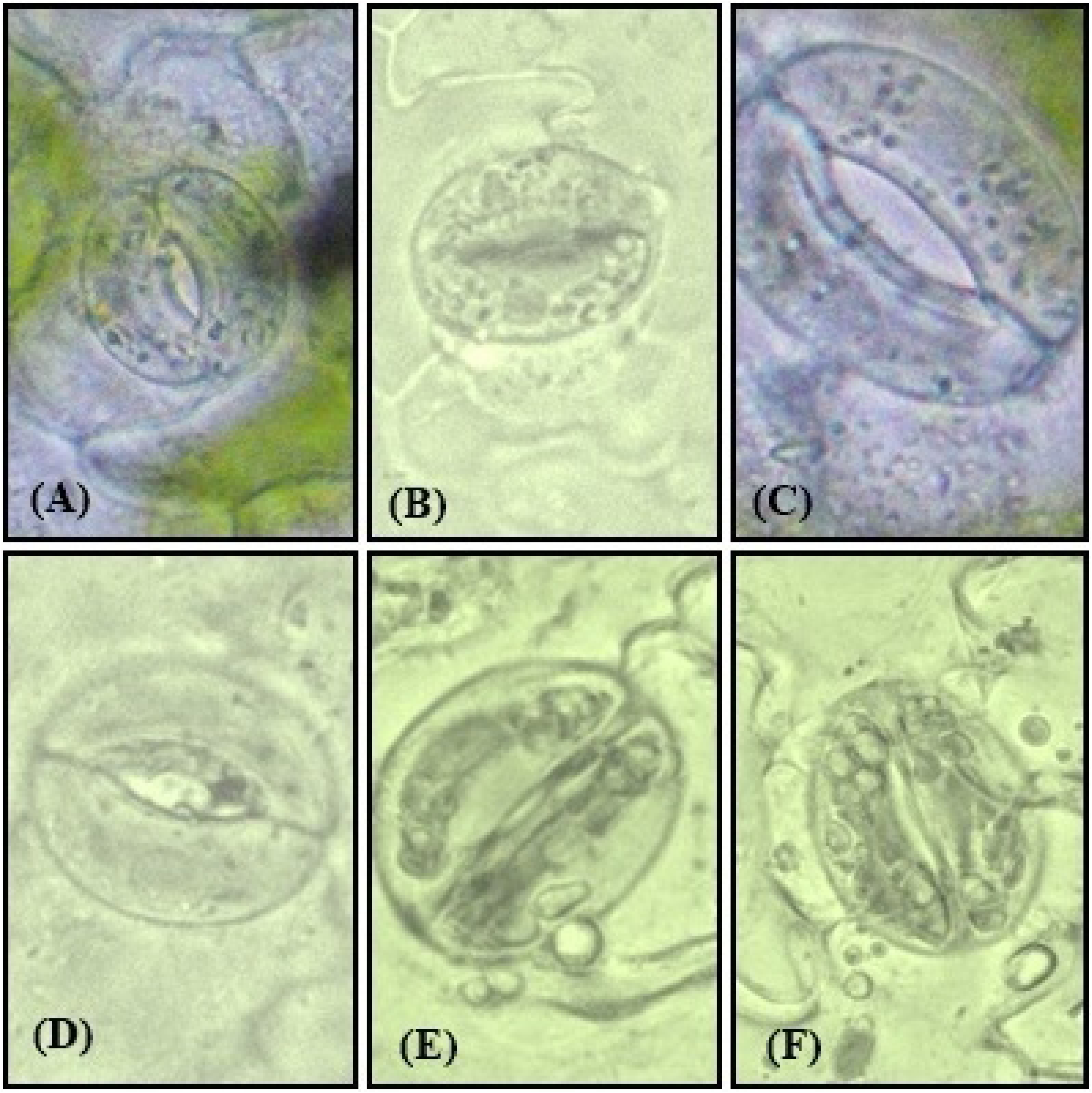
Stomatal response of *B. juncea* L. plants at 30 DAG under control **(A)**, Cu **(B)**, SNP **(C)**, SNP **+ Cu (D)**, FiCy+ Cu (**E)** and SNP **+** Cu +cPTIO **(F)** at 40× using compound microscope

### Effect of NO on NO generation, GSH content, plant dry mass, and leaf area

Figure 13, revealed that plants grown in Cu, exhibited increased NO generation by 4.3 times compared to control plants, but pre-germination supplementation of SNP decreased NO generation by 1.9 times compared to Cu exposed plants. However, minimum decrease i.e. not much significant in NO generation was noted by about 1.2 times with exogenously applied FiCy. Furthermore, application of cPTIO in combination with NO resulted in the reversal of the outcome of NO generation in Cu exposed plants.

**Fig 13.**
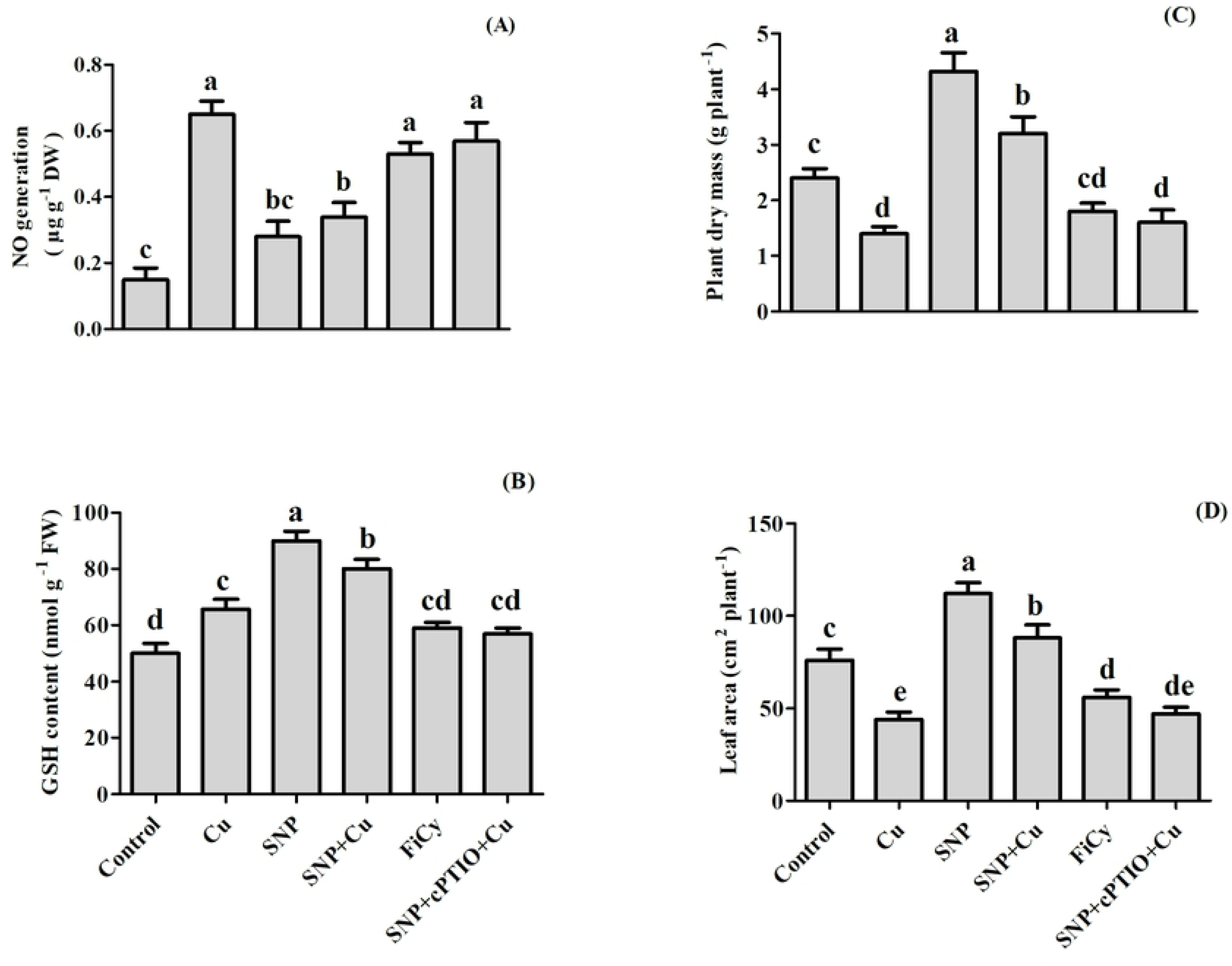
Effect of NO on NO Generation **(A),** GSH **(B)**, plant dry mass **(C)** and leaf area **(D)** of mustard (*B. juncea* L.) treated during pre-germination with water, 100 µM SNP or SNP with cPTIO and FiCy in presence or absence of 3 mM Cu at 30 DAG. Same letter above bars show that data did not differ significantly by LSD test at P < 0.05

Plants treated with 3.0 mM Cu increased GSH content while there was a reduction in plant dry mass and leaf area as compared to control. Plants with pre-germination treatment of SNP increased GSH content, and improved growth characteristics, plant dry mass and leaf area. Application of 100 µM SNP to Cu grown plants increased GSH content, plant dry mass and leaf area by 21%, 128%, and 100 %, respectively as compared to Cu treated plants, While the addition of cPTIO reversed the influence of exogenously applied NO and FiCy application showed no significant change to Cu exposed plants (Fig 13).

## Discussion

Copper is an essential micronutrient required for plant growth and development at low concentration, and it is a vital part of several biomolecules [6]. It has a high detrimental influence on seed germination and plant growth at high concentrations [4, 46]. NO as an important signaling molecule, has achieved an evident concern due to its crucial mitigating role in abiotic and biotic stresses in plants [38,47–49].

In the current study, SNP as an efficient NO donor was used, because it provides an absolute form of NO generation and supplies NO for an extensively longer duration as compared to other NO donors [50, 51]. However, SNP can also release cyanide (CN) and/or can form CN-Fe complex. It has been proposed that these complexes of CN may have an overlapping or distinct role with the role of NO alone on biological tissues [51]. Research evidence has shown that NO plays a significant role in seed germination of various species, such as warm-season grasses [30], *Arabidopsis* [29], and lettuce [27]. Furthermore, He et al. [52] provided evidence that NO attenuated the inhibition of germination of rice seed and ameliorated the inhibition of seedling growth caused by heavy metal like cafmium (Cd) in a concentration-dependent manner. They have also noticed that the protective effect of exogenous supplementation of SNP was reversed by addition of cPTIO, and FiCy had antagonistic effects in contrast to SNP. In agreement with the data published on other plants, our results proved the protective and stimulatory role of SNP on mustard seed germination. Conversely, the NO scavenger cPTIO reversed the NO-induced seed germination under Cu stress, while its analogue FiCy showed no change in germination percentage in comparison to Cu treated seeds. From our result, it came to the conclusion that NO derived from SNP but not any other compound FiCy was responsible for germination under Cu stress in mustard seeds. SNP pretreated mustard seeds were able to induce a rapid rise in the activity of amylase enzyme as compared to the control with a steady rise to a maximum till 24 hrs (Fig 6). According to Zhang et al. [53] the amylase activity was induced by NO was independent of gibberellic acid, showing that NO was involved in seed germination under various conditions.

Under Cu stress mustard seeds showed inhibition in germination and plants exhibited enhanced levels of TBARS and H_2_O_2_ content. These outcomes are in harmony with the earlier work of Hu et al. [31] and Fatma et al. [54], showing Cu and salt stress increased the accumulation of H_2_O_2_ and TBARS contents in wheat seeds and *B. juncea* leaves. Per et al. [38] emphasized that NO tends to decrease levels of TBARS and H_2_O_2_ content safeguarding the cell membrane to improve the cell membrane damage through lipid peroxidation. We further confirmed that the NO application reduced the production of ROS, H_2_O_2_ and O_2_^. –^ using DAB and NBT staining method in both Cu fed plants and germinating seeds. So far, the effect of NO on the inhibition of ROS accumulation of O_2_^.-^ and H_2_O_2_ using NBT and DAB staining in the germinating seeds has not been described in the literature. However, Fatma et al. [54] showed that NO alone or in combination with sulfur resulted in the reduction of ROS accumulation in salt-stressed plants. NO application also exhibited an inhibitory impact on Cu uptake in both shoots and roots in Cu fed plants (Fig 7 A, B). Our results are in accordance with the outcomes of Wang et al. [55] in tomato, Mostofa et al. [56] in rice and Zhang et al. [57] in tomato seedlings.

Our results also suggest that NO participates in enhancing the antioxidant enzyme activities, APX, GR, and SOD in pretreated germinating seeds and leaves of mustard grown under Cu stress. In compliance, various earlier published reports have proved the expression of antioxidant enzymes by the supplementation of NO in *Oryza sativa* under nickel toxicity [58], *Lycopersicon esculentum* under salinity stress [49], and in cotton under NaCl stress [59].

NO proved to be more promising and impactful in increasing photosynthetic characteristics, P_N_, Ci, gs, Rubisco activity, and PSII photochemical efficiency in presence and absence of Cu (Table 2). The mechanism behind the positive influence of NO on photosynthesis under Cu toxicity may be correlated to increased activity of antioxidant enzymes, protection of chlorophyll, and decreased ROS accumulation. It has been suggested that the NO can control the photosynthetic efficiency rate by controlling the size of stomatal aperture, hence influencing upon the stomatal conductance [54, 60]. SNP application also has some positive role in improving the activity of Rubisco enzyme and ultimately enhancing photosynthetic activity. The study is supported by numerous findings such as protection of chlorophyll damage and enhancement in photosynthetic pigments in NO-treated *Lolium perenne* under Cu stress [61], *Helianthus annuus* exposed to Cd stress [62], *Triticum aestivum* [63], *B. juncea* under salt stress [54]. Treatment of cucumber seedlings, with SNP enhanced the chlorophyll content, rate of photosynthesis, and transpiration rate and stomatal conductance [64]. These results were mainly due to the enhancement of antioxidant machinery, prevention, and recovery of chlorophyll damage and increased GSH content. In our study, the effect of NO prominently increased the content of GSH in the absence and presence of Cu stress. The study of Per et al. [38] also showed that NO accelerated GSH production in mustard plants treated with Cd.

SNP application ease the effect of Cu stress on plant dry weight and leaf area (Fig 12). Foliar spray of NO on several crops provided protection against Cu toxicity in *Lycopersicon esculentum* [57], *B. juncea* [65], *Nicotiana tabaccum* [66] and also alleviated effect of Ni stress in rice [58]. Growth improvement can also be associated with the escalation in antioxidant machinery after NO supplementation. The study of Bai et al. [67] in *L. perenne* revealed that the application of NO mitigated Pb stress, and alleviated negative effect on leaf growth by increasing the mineral nutrient and by opposing the oxidative stress parameters.

SNP application resulted in the reduction of NO generation in plants grown with or without Cu stress. However, minimum reduction of NO generation was found in Cu stressed plant and exogenously applied FiCy as NO analogue which did not release NO on its breakdown in Cu stressed plant. Further, application of cPTIO in combination with NO resulted in the reversal of the effect imposed by SNP on plants exposed to Cu. Wang et al. [68] also found that NO content increased in tomato plants in presence of Cu toxicity. This result also finds support from the study of Fatma et al. [54] in salt-stressed plants in which the NO alone in the combination of NO with S resulted in the maximum reduction of NO generation in the presence of salt.

## Conclusions

In conclusion, Cu induced toxicity resulted in a significant reduction of mustard seed germination, plant growth and photosynthetic efficiency. The findings of this study showed that NO pretreatment was able to improve seed germination by modulating antioxidant system, ROS accumulation, lipid peroxidation and amylase activity, these were the key elements in the defense mechanisms against Cu toxicity. The present research further proved that pretreatment of NO significantly mitigates Cu toxicity also during vegetative phase through improved antioxidant system, photosynthetic efficiency and reducing Cu induced accumulation of ROS accompanied by a reduction in lipid peroxidation of the mustard plant. These results suggest that pretreatment of NO could be employed as key biochemical approach for alleviating Cu-toxicity during seed germination and it not only results in improving germination rate but the applied NO also lead to improved vegetative phase of the plants. Hence pretreatment method is not only environment friendly but also cost effective.

## Acknowledgement

Authors are grateful to University Sophisticated Instruments Facility (USIF) of the Aligarh Muslim University for providing the necessary instrument facilities.

